# Spatial pattern and differential expression analysis with spatial transcriptomic data

**DOI:** 10.1101/2023.07.06.547967

**Authors:** Fei Qin, Xizhi Luo, Bo Cai, Feifei Xiao, Guoshuai Cai

## Abstract

The emergence of spatial transcriptomic technologies has opened new avenues to investigate gene activities while preserving the spatial context of tissues. Utilizing data generated by such technologies, the identification of spatially variable (SV) genes is an essential step in investigating tissue landscapes and biological processes. Particularly in typical experimental designs such as case-control or longitudinal studies, identifying SV genes between groups is crucial for discovering significant biomarkers or developing targeted therapies for diseases. However, current methods available for analyzing spatial transcriptomic data are still in their infancy, and none of the existing methods are capable of identifying SV genes between groups. To overcome this challenge, we developed SPADE for <u>s</u>patial <u>p</u>attern <u>a</u>nd <u>d</u>ifferential <u>e</u>xpression analysis to identify SV gene in spatial transcriptomic data. SPADE is based on a machine learning model of Gaussian process regression with a gene-specific Gaussian kernel, enabling the detection of SV genes both within and between groups. Through extensive simulations and real data analyses, we have demonstrated the superior performance of SPADE compared to existing methods in detecting SV genes within and between groups. The SPADE source code and documentation are publicly available at https://github.com/thecailab/SPADE.

## Introduction

Due to technological limitations, sequencing studies have overlooked valuable spatial information that is essential for understanding cell organization and interactions in space, as well as theirs links to tissue functions^1^. Excitingly, recent advancements in spatial transcriptomic technologies, such as 10x Visium^2^, Slide-seq^3^, Stereo-seq^4^, and smFISH^5^, enable gene-expression profiling with molecular and single-cell resolution while preserving spatial information of tissues^6^. These technologies provide key data for understanding disease mechanisms, especially in neurobiology and tumor biology, where biological activities are closely tied to the specific organization of cells^7^. For example, spatial transcriptomics of metastatic melanoma tumors have provided new insights into tumor progression and therapy outcomes^8^.

Spatially variable (SV) genes are critical for characterizing complex tissues using spatial transcriptomics^9^. These genes are defined as those with uneven, aggregated, or patterned spatial distribution of expression measures^9^, and they play a crucial role in biomarker discovery and drug development. However, current computational methods for identifying SV genes from spatial transcriptomic data are still in their early stages of development, and modeling transcriptomic data with numerous spatial locations remains statistically and computationally challenging^9^. SpatialDE^10^ decomposes expression variability into spatial and non-spatial components, and utilizes their ratio to detect SV genes based on the spatial variance. SPARK^9^ identifies SV genes using a linear spatial model with multiple Gaussian and periodic kernels. MERINGUR^11^ performs spatial autocorrelation and cross-correlation analysis to identify SV genes in spatial transcriptomic data. Despites these efforts to identify SV genes within a single group, there is currently no method available for detecting SV genes with differential spatial patterns between groups, which is essential for understanding changes in spatial patterns under different treatment conditions or time phases. Moreover, these existing methods use fixed hyperparameter values (i.e., the characteristic length scale) in their kernels to model how rapidly the covariance decays as a function of spatial distance, limiting their ability to capture specific spatial patterns, as shown in the results of this study.

To address these limitations, we developed SPADE for <u>s</u>patial <u>p</u>attern <u>a</u>nd <u>d</u>ifferential <u>e</u>xpression analysis to identify SV genes in complex tissues using spatial transcriptomic data. This method employes a Gaussian process regression (GPR) machine learning model^12^ with a gene-specific Gaussian kernel to enable accurate detection of SV genes. In addition to single-group pattern analysis, SPADE provides a framework for detecting SV genes between groups using a crossed likelihood-ratio test. To evaluate the performance of SPADE, we benchmarked it against existing spatial pattern investigation methods (i.e., SpatialDE^10^, SPARK^9^, MERINGUE^11^) using extensive simulation studies and real data analyses with pre-identified SV genes. We also assessed the performance of SPADE in detecting SV genes between groups using simulations and a real dataset across the regeneration stages of axolotl telencephalon. The results demonstrated that SPADE outperforms other methods in identifying SV genes both within and between groups. Therefore, this study provides a powerful method for identifying SV genes in spatial transcriptomic data, with the potential to improve biomarker discovery and drug development.

## Results

### The framework of SPADE

To enable the identification of SV genes from spatial transcriptomic data, both within a single group and between groups, we developed the SPADE method based on a GPR model^12^ with a Gaussian kernel. The framework of the SPADE method is illustrated in Fig. 1. Briefly, the original read count data are first normalized into continuous data using a two-step normalization strategy. The optimal length-scale hyperparameter in the kernel for each gene is further estimated. Subsequently, a quadratic Q statistic is constructed to compute the *P-*value for detecting SV genes within groups. In addition, SPADE utilizes a crossed likelihood-ratio test to detect SV genes between groups. The performance of SPADE was assessed through extensive simulation studies and real data analyses, for the identification of SV genes both within groups and between different groups.

**Figure 1:**
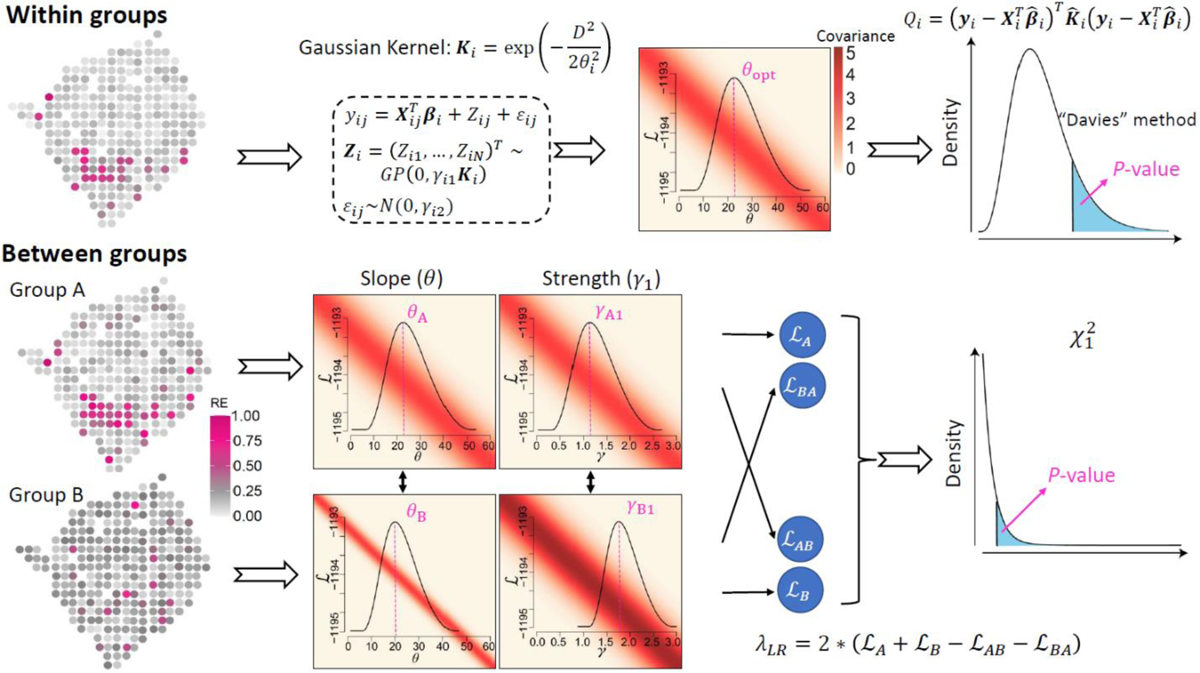
Framework of SPADE. SPADE was developed based on a Gaussian process regression model with a Gaussian kernel to identify spatially variable (SV) genes within groups and between groups. The optimal length-scale hyperparameter (θ) in the kernel function was estimated for each gene and the *P*-value was calculated based on a quadratic *Q* statistic with the Davies method. To detect SV genes between groups, SPADE exchanged the optimal parameters (θ, γ_1_) estimated from two groups, and calculated log likelihoods (ℒ) for themselves (i.e., ℒ_A_, ℒ_B_) and their counterparts (i.e., ℒ_AB_, ℒ_BA_). θ controls the slope of covariance to decay and γ_1_ controls the strength of covariance. Then a crossed likelihood-ratio test was utilized to calculate the *P*-values.

### Evaluation of SPADE to identify spatially variable genes within a group

#### Simulated patterns

We first assessed the performance of SPADE to identify SV genes within groups using two sets of simulations and three real datasets (SeqFISH, MERFISH, MOB). SPADE was evaluated by simulation and real data studies in identifying spatial patterns, and compared to existing methods (SpatialDE, SPARK and MERINGUE) using *F*1 scores. In the simulations involving hotspot and streak patterns, 200 genes and 200 measurement spots were simulated by mimicking the MOB dataset, with varied proportions of marked spots (10%, 20%, 30%) and different expression strength in fold changes (1.2, 1.3, 1.4, 1.5, 1.6). For both hotspot and streak patterns, SPADE consistently achieved the highest *F*1 score among all methods in almost all scenarios (Fig. 2), followed by SPARK, SpatialDE, and MERINGUE. For instance, in the hotspot simulation pattern with a fold change of 1.4 for 20% marked spots, SPADE achieved a higher *F*1 score of 0.83 to identify SV genes, than other existing methods (SPARK: 0.75, SpatialDE: 0.51, MERINGUE: 0.04) (Fig. 2A). The outperformance of SPADE was also observed in detecting streak patterns (Fig. 2B). When the proportion of marked spots or the difference in gene expression increased, the spatial patterns were more evident, and all methods showed an improved performance on pattern detection.

**Figure 2:**
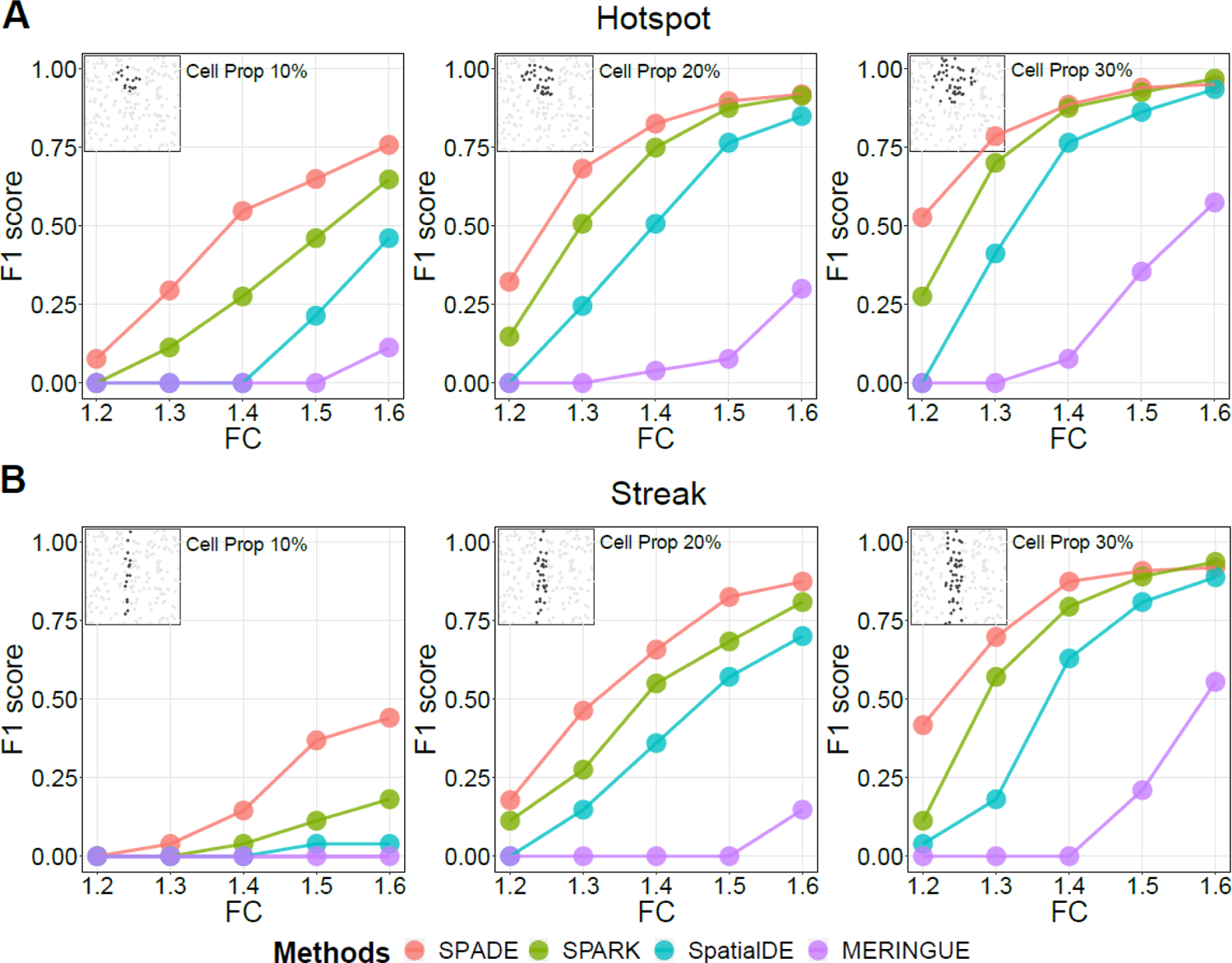
Assessment of SPADE to identify spatially variable genes within groups using simulations. Simulations were utilized to evaluate the performance of SPADE to identify spatially variable genes within groups. SPADE was compared to existing methods with *F*1 scores including SpatialDE, SPARK and MERINGUE. Two distinct spatial patterns were included in the simulations, including the (A) hotspot and (B) streak patterns. Different proportions (10%, 20%, 30%) of marked spots were considered in each pattern. We also utilized various expression differences in fold changes (1.2, 1.3, 1.4, 1.5, 1.6) for the marked spots to evaluate how signal strength affects the performance of different methods.

#### Pattern from real data in simulations

We further evaluated all the methods in detecting spatial patterns observed in real datasets (SeqFISH, MERFISH, MOB). SPADE was compared to existing methods using simulated expression of various fold changes (1.5, 2.0, 2.5, 3.0, 3.5) in marked spots for three distinct patterns from real datasets. SPADE achieved the highest *F*1 score in almost all the scenarios (Fig. 3). For example, in the SeqFISH dataset simulation with a fold change of 3.0 (Fig. 3A), SPADE obtained the highest *F*1 score of 0.67, surpassing SPARK (0.43), SpatailDE (0.28), and MERINGUE (0.50). In the MERFISH dataset with a large number of spots (n=4,975) (Fig. 3B), SPADE showed the best performance that consistently achieved an *F*1 score close to 1.0 across different fold changes. The performance of SPARK and SpatialDE dramatically declined in detecting SV patterns with a larger difference in gene expression, characterized by sparser marker spots. Notably, a higher variation in the estimated length-scale hyperparameter values was observed when the expression difference in fold change increased (Supplementary Figure S1). This finding indicated that existing methods with fixed hyperparameters were not optimized to accurately model all genes and could consequently be powerless to identify more complex spatial patterns (e.g., sparse marker spots). Increased detection power was observed for all methods when identifying the spatial pattern with larger fold changes in the MOB dataset (Fig. 3C), consistent with the finding for detecting simulated hotspot and streak patterns discussed above and shown in Fig. 2.

**Figure 3:**
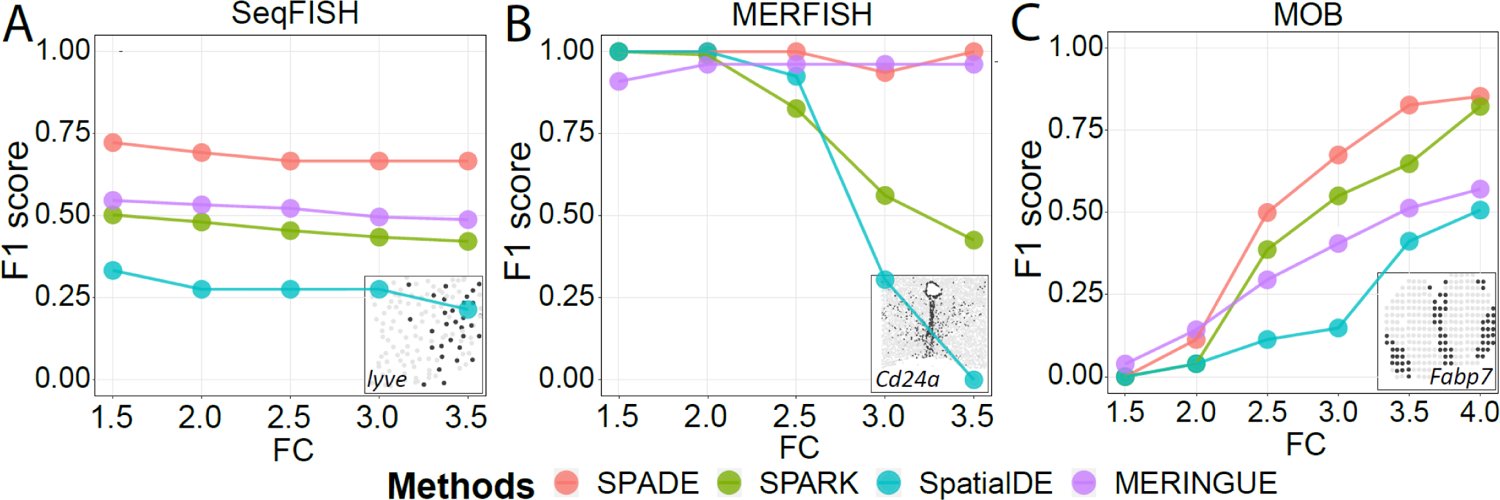
Assessment of SPADE to identify spatially variable genes within groups using real data-based simulations. Simulations with significant spatial patterns from real datasets, including (A) SeqFISH, (B) MERFISH and (C) MOB, were utilized for the evaluation of SPADE to identify spatially variable genes. SPADE was compared to existing methods with *F*1 scores including SpatialDE, SPARK and MERINGUE. Based on these predefined differential spatial patterns, spots with expression value > 0.75 quantile of all spots were assigned as marked spots in simulation data, including *lvye* in the SeqFISH dataset, *Cd24a* in the MERFISH dataset, and *Fabp7* in the MOB dataset. These marked spots were given various expression differences in fold changes (1.5, 2.0, 2.5, 3.0, 3.5).

#### Real data analyses

Furthermore, with real data analyses, we evaluated the performance of SPADE in identifying within-group SV genes using pre-identified “true” marker genes from the SeqFISH and MOB datasets (see Methods). SPADE was assessed using *F*1 scores and compared to existing methods. On the SeqFISH dataset, SPADE outperformed other methods to identify SV genes (e.g., *lyve*, Fig. 4A top) with an *F*1 score of 0.63, compared to SPARK (0.54), SpatialDE (0.48), and MERINGUE (0.29) (Fig. 4A bottom). On the MOB dataset, SPADE identified a smaller number of SV genes (n=150, e.g., *Fabp7*, *Pcp4*, Fig. 4B top) as compared to SPARK (n=772) and MERINGUE (n=3,918) (Fig. 4B bottom). SpatialDE did not identify any SV genes. Notably, SPADE showed a higher enrichment (50.0%) in cell type-specific marker genes, comparing to those from SPARK (33.7%) and MERINGUE (12.5%). Additionally, we used highlight genes with enriched expression in the mitral cell layer (n=10) from MOB original study to evaluate the performance of all methods and found a higher “true positive” rate of SPADE (1.33%) than SPARK (1.04%) and MERINGUE (0.20%).

**Figure 4:**
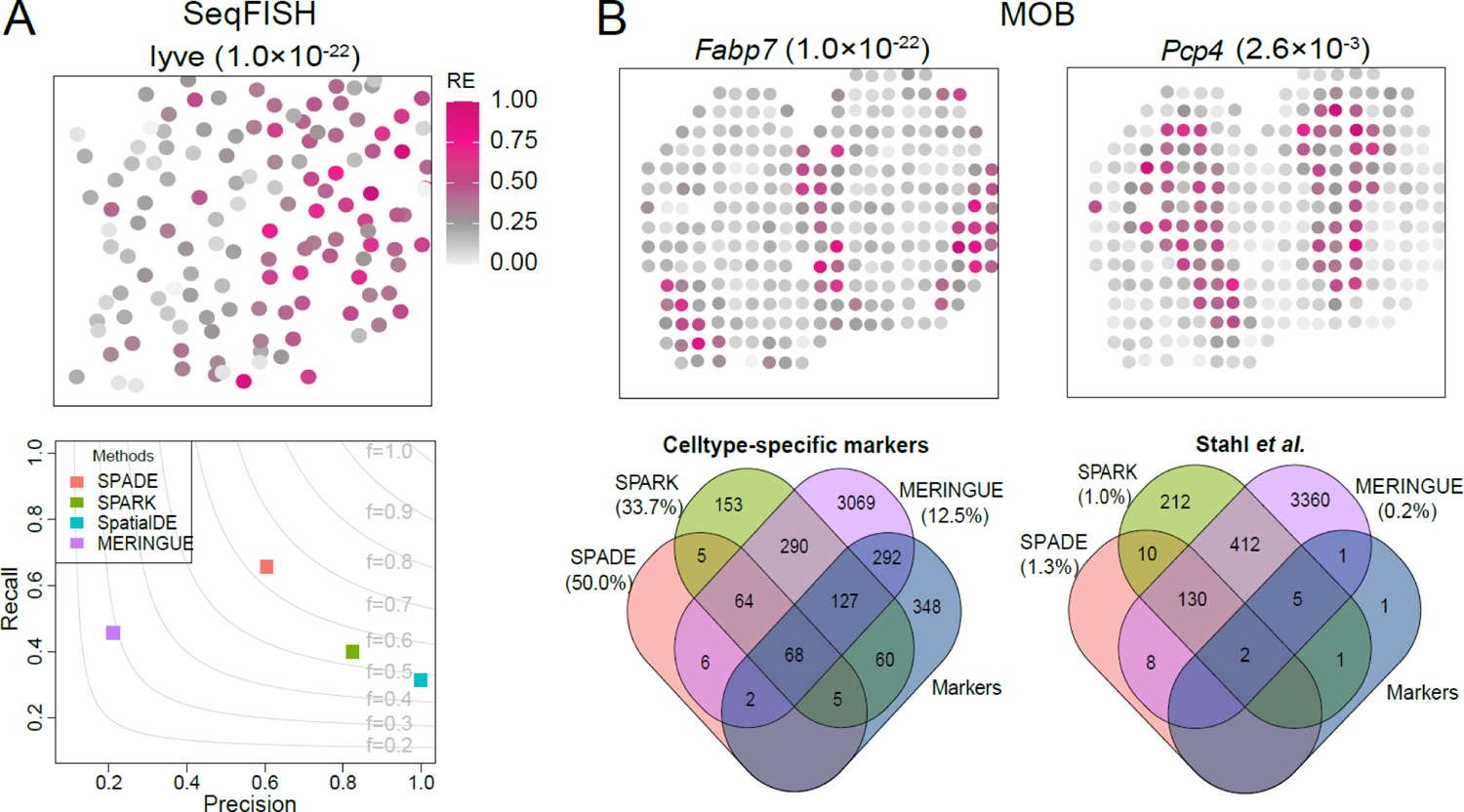
Assessment of SPADE to identify spatially variable genes within groups using real data analyses. Real datasets, including (A) SeqFISH and (B) MOB, with predefined “true” markers were used for the evaluation of SPADE to identify spatially variable (SV) genes within groups. SPADE was compared to existing methods including SpatialDE, SPARK and MERINGUE. In the seqFISH dataset, 35 genes were pre-selected from its original study as cell identity markers for the assessment of SPADE. In the MOB dataset, two different lines of predefined “gold standard” were used to validate the SV genes identified from each method, including 902 cell type-specific marker genes from a recent single-cell RNA sequencing study and 10 highlighted marker genes in the olfactory system. Representative SV genes identified by SPADE in each dataset are shown, with *P* values from SPADE inside parentheses.

In summary, with two sets of simulations and real data analyses, SPADE was found to be more accurate than existing methods in detecting within-group SV genes in spatial transcriptomic data.

### Evaluation of SPADE to identify spatially variable genes between groups

#### Simulated patterns

With both simulations and real data analyses, we also evaluated the performance of SPADE in detecting SV genes between groups. Mimicking the MOB dataset, simulation data with 200 genes and 200 spatial spots were generated in two groups, with and without spatial patterns respectively. Various scenarios were simulated with different marker proportions (10%, 20%, 30%), expression differences in fold changes (1.5, 2.0, 2.5, 3.0, 3.5), and signal shapes (hotspot, streak). The receiver operating characteristic (ROC) curve was generated and the areas under the ROC curve (AUC) were utilized to assess the power of different methods with pre-defined markers. No existing method was developed specifically for this purpose but methods for differential expression analysis provide a possible option. Therefore, SPADE was benchmarked against widely used differential expression analysis methods, including edgeR, DESeq2, limma-voom, and MAST. In almost all scenarios, SPADE outperformed other methods, and all existing differential expression analysis methods in study showed no power in identifying SV genes between groups in spatial transcriptomic data (Fig. 5A-B). Consistent with the detection of within-group SV genes, SPADE demonstrated higher power in detecting between-group SV genes with a higher proportion of marked spots or a larger difference in gene expression.

**Figure 5:**
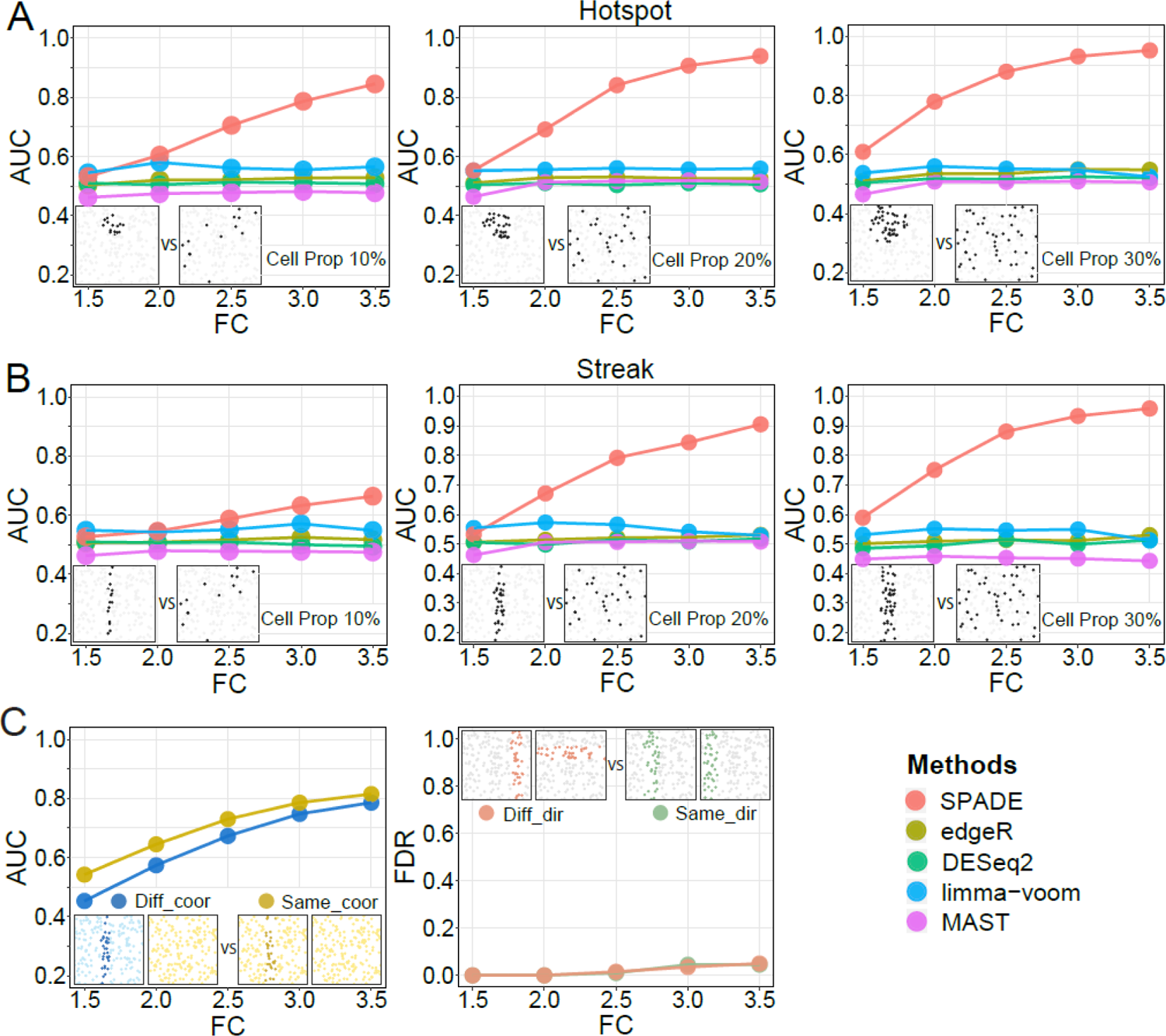
Assessment of SPADE to identify spatially variable genes between groups using simulations. SPADE was compared to typical differential expression analysis methods including edgeR, DESeq2, limma-voom and MAST. Two groups of data were generated with one group assigned with the (A) hotspot or (B) streak patterns, considering various marked spots proportions (10%, 20%, 30%) and expression differences in fold changes (1.5, 2.0, 2.5, 3.0, 3.5). We also evaluated the effects of spot coordinates and pattern directions to the performance of SPADE with the streak patterns and marked spots proportion of 20% (C). The areas under the receiver operating characteristic curve (AUC) were utilized to assess the power, and the false discovery rate (FDR) was used to assess the type I error of SPADE, respectively. Diff_coor: different coordinates; Same_coor: same coordinates; Diff_dir: different pattern directions; Same_dir: same pattern directions.

#### Robustness to unaligned quantifications

There is challenge of detecting SV genes between unaligned quantifications, such as those with different spots coordinates, different subjects, or angles (e.g., directions of the streak pattern). SPADE implemented a crossed likelihood-ratio test to address this challenge. In our simulation study with 20% marker proportion, SPADE showed a close AUC in detecting steak patterns between groups with different spot locations compared to those with the same locations (Fig. 5C left). Moreover, similarly small FDRs were given by SPADE in groups with different signal directions compared to those with the same signal directions (Fig. 5C right). A similar performance was also observed in detecting the hotspot pattern (Supplementary Figure S2) and patterns with 10% and 30% marker proportions (Supplementary Figure S3-S4). These findings collectively indicate that SPADE is robust to the variation of pattern directions and locations.

#### Real data analyses

The performance of SPADE to detect SV genes between groups was further evaluated using a real dataset of ARRISTA with two different post injury states (i.e., 2DPI, 5DPI) from brain tissue of axolotl. Among a total of 2,020 genes, 481 SV genes (e.g., *CDH1D*, *PDE2A*, Fig. 6A) were identified by SPADE between these two states. These identified SV genes were significantly (*P*-value < 2.2 × 10^−1^^6^) overlapped with cell type specific markers from the original study (Fig. 6B). This indicated that SPADE identified SV genes related to cell types which were crucial for the development of brain tissue in axolotl. These SV genes from SPADE were enriched in pathways related to tissue regeneration related biological processes, including organ development, carbohydrate response, oxidative stress, mitochondrial activity, metabolic process, and wound healing, which were highly involved in tissue regeneration (Fig. 6C). This well aligned with reports that oxidative stress has been found to play important roles in tissue regeneration of various species^13, 14^; metabolism regulates cell proliferation and regeneration^15^; mitochondria and carbohydrates play crucial roles in providing the necessary energy for tissue repair and regeneration^16, 17^; SV genes identified were found with roles in wound healing and neuron activity, potentially contributing to axolotl brain regeneration post injury^18, 19^.

**Figure 6:**
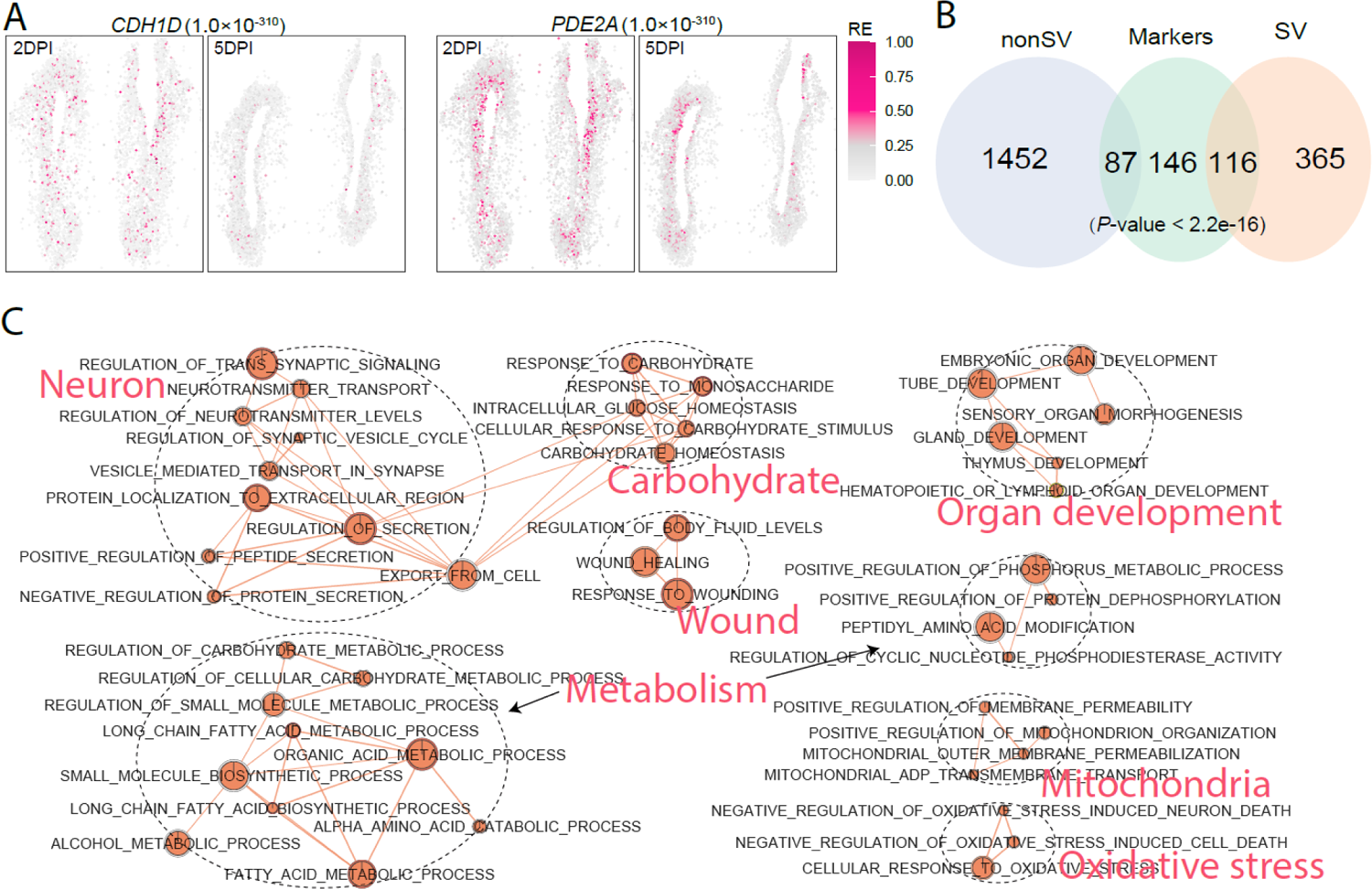
Assessment of SPADE to identify spatially variable genes between groups using real data analyses. We evaluated the performance of SPADE to identify spatially variable (SV) genes between groups using a real spatial transcriptomic dataset with axolotl telencephalon (ARRISTA). Two different post injury stages (i.e., 2DPI, 5DPI) were included in the analysis. SV genes between these two groups were identified using the SPADE method and then compared to pre-selected cell type identity markers from its original study. (A) Representative SV genes identified by SPADE are shown, with *P* values from SPADE inside parentheses. (B) Pearson’s chi-square test was utilized for the evaluation of the association between SV genes from SPADE and those “true” markers. (C) The pathway enrichment analysis was conducted based on these SV genes with connection networks generated using the Enrichment Map implemented in Cytoscape. Node size corresponds to the number of genes within the pathway. Edge weight corresponds to the number of genes found in both connected pathways.

## Discussion

The emerging of spatial transcriptomics has greatly advanced research in various biomedical domains^20, 21^. It provides an opportunity to identify genes that exhibit spatial expression patterns but requires specialized and sophisticated methods. In this study, we developed the SPADE method to address this demand. Through extensive simulations and real data analyses, we demonstrated the desirable performance of SPADE in detecting SV genes from spatial transcriptomic data. One merit of SPADE resides in its gene-specific length-scale hyperparameter within the kernel function. This feature contributes significantly to the superior performance of SPADE in identifying spatial patterns compared to existing methods, which typically utilize length-scale-fixed kernels. Additionally, SPADE for the first time enables detection of SV genes between groups, making it suitable for experimental designs of different treatment conditions or time phases.

The comparison of two spatial patterns poses challenges due to inconsistencies in spots coordinates and measuring angles. SPADE is robust to those inconsistencies by its crossed likelihood-ratio test strategy. Our study showed that conventional differential expression methods are inadequate for detecting SV genes with distinct spatial patterns in two groups. This is not surprising as these conventional differential expression methods test difference in mean expression but not spatial patterns. Applying SPADE, we successfully identified SV genes related to cell types between two different post-injury states of the axolotl brain. We believe SPADE has a broad application scope to study spatially variable patterns between different developmental stages, which has been widely studied to elucidate the spatial dynamics of embryonic heart development^22^, mammalian spermatogenesis^23^, intestinal development^21^ and human endometrium^24^.

As SPADE estimates parameters for each gene, computational time can be challenging for modeling large dataset with spatial information (e.g., > 5,000 spots). For computational speed (Supplementary Table S1), with 249 genes and 131 spots in the SeqFISH dataset, the time required for SPADE to detect SV genes was 1.16 mins, which was slightly longer than that of SPARK (1.08 mins) and MERINGUE (0.01 mins) but shorter than that of SpatialDE (1.68 mins) at a desktop workstation with an Intel Core i5 CPU 2.6 GHz processor and 8.0 GB of RAM. Nevertheless, given the improved performance of SPADE to identify SV genes, we don’t consider the computational time to be a significant issue for small/medium datasets. For large datasets, advanced computational infrastructure such as high-performance computing could be utilized to largely alleviate this issue.

In general, our SPADE method has provided a valuable tool for the investigation of differential expression patterns in spatial transcriptomic data. It also has the potential to be extended by incorporating other tissue information (e.g., histopathological images, temporal information) into the model for more accurate inference of significant gene markers. For example, histopathological images provide high-resolution cellular information, which will be helpful to enhance the spatial expression patterns^25^; Incorporating temporal information can contribute to the discovery of spatial expression patterns associated with dynamic changes of cell states and types^26^. These warrant future studies.

## Methods

SPADE was developed based on a GPR model with a Gaussian kernel to capture the relationship between gene expression and other covariates (i.e., cell groups) incorporating the spatial information of cells. Various technologies possess distinct measurement resolutions, ranging from sub-cellular to sets of cells located in small regions. For consistency, we will use “spot” as a general detection region to describe the model across the manuscript.

### Data normalization

SPADE took read counts as input and transformed them into continuous data using a two-step normalization strategy to achieve efficient estimation of parameters. First, the mean-variance dependency in sequencing data was stabilized using Anscombe’s transformation with R^′^_i_ = log (R_i_ + 1/Φ)^27^, where R_i_represented the read counts for i-th gene, and Φ was the common overdispersion parameter^28^ used to capture the dependency of mean and variance across all genes with *Var(R_i_) = E(R_i_) + Φ · E(R_i_)^2^*. Next, the effects of library size were regressed out from the transformed data R^′^ to generate the normalized continuous data^10^, which were utilized in modeling of the SPADE method as shown below.

### Notation and models

Let **Y** = (y_ij_)_P×N_ denote the normalized continuous expression data for the i-th (i = 1, …, P) gene and the j-th (j = 1, …, N) spatial spot, with two-dimensional spatial coordinates **S**_j_ = (S_j1_, S_j2_). For each gene, a GPR model^12^ is constructed to model the spatial pattern of gene expression data that

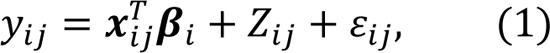

where x_ij_is a *k*-covariate vector that includes *k* −1 covariates (e.g., batch effect, cell type) and a scalar one showing the intercept for the mean log-expression of the gene across all spots. β_i_ is the vector of coefficients for these covariates, and ε_ij_ is the residual error that is independently and identically distributed from N(0, γ_i2_). *Z*_ij_is a zero-mean stationary Gaussian process modeling the spatial correlation pattern with Z_i_ = (*Z*_i1_, …, *Z*_iN_)^T^∼ MVN(0, γ_i1_K_i_), where K_i_ is a covariance kernel function of a multivariate normal distribution for the i-th gene based on pairwise distances of all spots, and γ_i1_is a scaling factor. Kernel functions are widely used to map data with spatial information into high-order feature space and capture the nonlinear interactions among spots^29^. In SPADE, a Gaussian kernel is used to model the covariance matrix among spots, denoted as 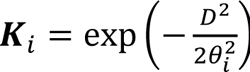, where D = ‖**S** − **S**^′^‖ is the Euclidean distance matrix^30^ of spots based on spatial coordinates. θ_i_in the kernel function is the hyperparameter of length-scale to model the curvature of the covariance to the distance between spots, where a larger θ_i_ corresponds to a smoother covariance change^12^. Thus, the total covariance of y_i_ = (y_i1_, …, y_iN_)^T^ is Σ_i_ = γ_i1_K_i_ + γ_i2_*I*, where *I* is an *n*-dimensional identity matrix. γ_i1_effectively measures the expression variance captured by spatial patterns, and γ_i2_measures the expression variance owing to random noise. As far, the problem of identifying SV genes has been transferred into testing the null hypothesis “H_0_: γ_i1_ = 0”. The power of SV gene identification relies on how effectively spatial kernel matrix K captures the significant spatial patterns exhibited by genes of our interest. Our goal is to estimate Θ_i_ = (β_i_, θ_i_, γ_i1_, γ_i2_) for the i-th gene.

### Parameter estimation and identification of spatially variable genes

After marginalizing over function values Z_i_, we obtain the multivariate normally distributed marginal likelihood of the expression values p(y_i_|y_i_, Θ_i_) = MVN(y_i_|y^T^β_i_, Σ_i_), where y_i_ = (x_i1_, …, x_iN_)^T^. p(y_i_|y_i_, Θ_i_) is calculated efficiently through spectral decomposition^31^ of the covariance matrix K_i_. To estimate Θ_i_, we optimize the log marginal likelihood with respect to the coefficients of covariates β_i_, kernel hyperparameter θ_i_, spatial variance factor γ_i1_, and random variance γ_i2_ using an optimization strategy of golden section search^32^. More details on the estimation of these parameters are provided in the Supplementary Methods.

After selecting the optimal length-scale hyperparameter (^θ^^_i_) for the i-th gene, under the null hypothesis (γ_i1_ = 0), we estimate the coefficients for covariates (^β^^_i_) using a score-based variance-component test, SKAT^33^. A *Q* score statistic can be easily calculated based on the GPR model with a quadratic form 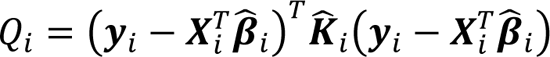^34^, where ^K^^ is the estimate of *Q*_i_ = (y_i_ − y_i_ β_i_) K_i_(y_i_ − y_i_ β_i_) kernel covariance matrix based on the optimal length-scale hyperparameter ^θ^^_i_. Theoretically, *Q*_i_follows a mixture of independent chi-square distributions with mixing weights that depend on the eigenvalues of the kernel matrix. Based on this, the Davies method^35^ is utilized to compute a P-value for the i-th gene. P-values across all genes were adjusted with the Benjamini and Hochberg (BH) method^36^ to correct for multiple testing and control for false discovery.

Furthermore, the SPADE method can identify SV genes between groups (e.g., group A and B) based on a crossed likelihood-ratio test with spatial transcriptomic data. To achieve this, we first estimate the optimal hyperparameter in the kernel function for each group, respectively. Thus, for each gene, the log likelihood (ℒ) in group A and B can be easily calculated with its optimal kernel matrix, referred to as ℒ_A_ and ℒ_B_, separately. Then we use the optimal parameter values (γ_i1_, θ_i_) estimated from each group to compute alternative log likelihoods for their counterparts (ℒ_AB_, ℒ_BA_). By comparing alternative likelihoods to optimal log likelihoods, we can test whether these two groups have the same spatial patterns. The likelihood-ratio test statistic is computed with λ_LR_ = 2 ∗ (ℒ_A_ + ℒ_B_ − ℒ_AB_ − ℒ_BA_) and *F* test with degree freedom of one is used to calculate P-values. Also, P-values across all genes were adjusted using the BH method^36^.

### Datasets

Four spatial transcriptomic datasets, including MOB^20^, SeqFISH^37^, MERFISH^38^ and ARTISTA^39^, were analyzed in our study to evaluate the performance of SPADE in identifying SV genes. The MOB, SeqFISH and MERFISH datasets were utilized for evaluating SPADE to detect SV genes within groups, while the ARTISTA dataset was used to evaluate the performance of SPADE in identifying SV genes between groups.

The MOB dataset^20^ comprises spatial transcriptomic data with mouse olfactory bulb (MOB) measured through spatial barcoding. Raw read counts were downloaded from Spatial Transcriptomics Research (http://www.spatialtranscriptomicsresearch.org). In this study, we analyzed the “MOB Replicate 11” sample which was also used by the study of SPARK^9^. We filtered out genes expressed in less than 10% spots and spots with less than 10 total read counts. This filtering process resulted in a final set of 11,274 genes on 260 spots for simulation studies and real data analyses.

The SeqFISH dataset^37^ was collected on the mouse hippocampus with 249 genes measured on 257 spots by smFISH. 35 out of these 249 genes were pre-identified in its original study as cell identity markers. Raw read counts with the field 43 dataset were obtained from its original report. Following the Trendsceek study^40^, we filtered out cells located at the boundary to relieve border artifacts, resulting in a final set of 249 genes measured on 131 spots for evaluating SPADE to identify SV genes within groups in both simulations and real data analyses.

The MERFISH dataset^38^ was downloaded from Dryad platform with the preoptic region of the hypothalamus in mouse measured by smFISH. We used slice from animal 18 for analysis, which was used in the study of the study of SPARK^9^ and included 160 gene and a large number (>5,000) of spot. After removing the ambiguous cells identified as putative doublets following the SPARK study^9^, we retained a final set of 160 genes on 4,975 spots for evaluating SPADE to identify SV genes within groups.

The ARRISTA dataset^39^ provided the landscape of gene expression across the regeneration stages of axolotl telencephalon at single cell resolution by Stereo-seq, which was downloaded from https://db.cngb.org/stomics/artista/. Spatial transcriptomic data for two (2DPI) and five (5DPI) days post-injury brain tissue of axolotl were analyzed and SV genes between these groups were identified. After filtering out less informative genes expressed in less than 20% spots and spots with less than 1,000 total read counts, 2,168 overlapped genes in both groups (3,961 spots in the 2DPI group and 3,128 spots in the 5DPI group) were included in the analysis. 349 cell type markers pre-identified in its original study^39^ were utilized for performance evaluation of SPADE to identify between-group SV genes.

### Evaluation of SPADE using simulations

Simulation studies were conducted to evaluate the performance of SPADE to identify SV genes both within and between groups. In particular, SPADE was benchmarked to existing methods including SpatialDE^10^, SPARK^9^ and MERINGUE^11^ to assess its ability to detect within-group SV genes. In simulations with two groups, SPADE was benchmarked to existing methods which were developed for typical differential expression analysis in RNA-seq data including edgeR^41^, DESeq2^42^, limma-voom^43^ and MAST^44^.

### Within-group spatially variable gene simulation

Simulations were implemented in two sets for the evaluation of SPADE to detect SV genes within groups. In the first set, simulation studies mimicked the MOB dataset, as introduced in the section of Datasets, to generate simulation data by modifying single-cell RNA-seq data simulator *splatter*^45^. We generated 200 genes and 200 spots with spots coordinates simulated with a random-point-pattern Poisson process^46^. These simulation data were treated as “pattern free” data of which 50 genes were randomly picked to assign different spatial patterns as “true” SV genes for the evaluation of SPADE. Two distinct spatial patterns were included in the simulations, including the hotspot and streak patterns, and different proportions (10%, 20%, 30%) of marked spots were considered. Marked spots provide distinct expression strength compared to other spots. Moreover, to evaluate how signal strength difference affects the performance of SPADE, we utilized various expression differences in fold changes (1.2, 1.3, 1.4, 1.5, 1.6) for these marked spots.

In the second set of simulations with more complicated patterns, we simulated gene expression data with spatial patterns observed in real datasets from the SeqFISH, MERFISH and MOB datasets, respectively. A general simulation strategy was utilized to generate “pattern free” data based on a negative binomial distribution NB(10, 0.1). Representative genes from real datasets were then mimicked to construct spatially variable patterns in simulation studies. The SV genes identified by all analysis methods were defined as representative genes in simulations, including *lvye*, *Cd24a* and *Fabp7* in the SeqFISH, MERFISH and MOB datasets, separately. Based on these representative genes, spots with expression value > 0.75 quantile of all spots were assigned as potential marked spots and given various expression differences in fold changes (1.5, 2.0, 2.5, 3.0, 3.5). In simulations, we generated data with 200 genes and the same number of spots for each dataset. Still, 50 genes were randomly selected as “truth” SV genes for the assessment of SPADE. In both sets of simulations, the performance of the SPADE method to detect SV genes was assessed by *F*1 score 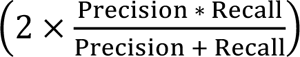 with the precision rate defined as 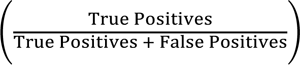 and recall rate defined as 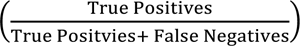.

### Between-group spatially variable gene simulation

The MOB dataset was used to generate two groups of simulation data as “pattern free” data with 200 spots each and 200 genes in total. In the first group, 50 genes were randomly selected as “truth” SV genes and assigned with different spatial patterns (i.e., hotspot and streak). We also considered various proportions (10%, 20%, 30%) and signal differences in fold changes (1.5, 2.0, 2.5, 3.0, 3.5) for these marked spots. AUC was utilized to assess the power of different methods.

Two patterns could be identified as non-SV patterns with similar signal shapes and strength but from different biopsy specimens with different locations or angles. Therefore, we assessed the robustness of SPADE in detecting SV genes between groups using data with different spots coordinates or pattern directions. Specifically, we compared the detection AUC of SPADE in two groups with the same locations to those with different locations. We also benchmarked SPADE in groups with the same pattern angles to those with different angles. Thus, false discovery rate (FDR) was used to assess the type I error of SPADE. Simulation studies included both hotpots and streak patterns, with different proportions (10%, 20%, 30%) of marked spots and various signal strength with fold changes of 1.5, 2.0, 2.5, 3.0, and 3.5.

### Evaluation of SPADE using real data analyses

#### Within-group spatially variable gene analyses

We assessed the performance of SPADE to detect SV genes within groups using real datasets, including the SeqFISH and MOB datasets (details are shown in the Datasets). In the SeqFISH dataset, 35 genes were pre-selected from its original study as cell identity markers which were utilized as “true” SV genes for comparing SPADE to existing methods (SpatialDE^10^, SPARK^9^ and MERINGUE^11^). In the MOB dataset, two independent lines of evidence were used to validate the SV genes identified from each method. First, 10 marker genes with highly enriched expression in the olfactory system from its original study^20^ were used as “true” markers. Second, 902 cell type-specific marker genes from a recent single-cell RNA sequencing study^47^ in the olfactory bulb were utilized as “gold standard” for the assessment of different methods. Similarly, all methods were evaluated using precision rate, recall rate and *F*1 score.

#### Between-group spatially variable gene analyses

We further evaluated the performance of SPADE to identify SV genes between groups using a real spatial transcriptomic dataset (i.e., ARRISTA) with axolotl telencephalon, which was described above. With two groups of data from different post injury stages (i.e., 2DPI, 5DPI) and 2,020 consistent genes, SV genes between these two groups were identified using the SPADE method and compared to pre-selected cell type identity markers from its original study^39^. Pearson’s chi-square test was utilized to investigate the association between SV genes from SPADE and the “true” markers. The Gene Ontology (GO) biological process pathway enrichment analysis^48^ was conducted based on these significant genes. Further, the summary statistics from pathway analysis were used to generate connection networks using the Enrichment Map implemented in Cytoscape^49^.

## Data availability

The MOB dataset was downloaded from Spatial Transcriptomics Research (http://www.spatialtranscriptomicsresearch.org). The SeqFISH dataset was downloaded from the supplementary file of the original paper (https://www.cell.com/cms/10.1016/j.neuron.2016.10.001/attachment/759be4dc-04a6-4a58-b6f6-9b52be2802db/mmc6.xlsx). The MERFISH dataset was available in the Dryad platform with mouse preoptic region of the hypothalamus (doi: https://datadryad.org/stash/dataset/10.5061/dryad.8t8s248). The ARRISTA dataset can be explored and downloaded from https://db.cngb.org/stomics/artista/.

## Code availability

SPADE source code and documentation are publicly available at https://github.com/thecailab/SPADE.

## Author contributions

F.Q, X.L, B.C and G.C developed the method. F.Q and G.C performed simulations and analyzed real data. F.Q, F.X and G.C wrote the manuscript.

## Competing interests

The authors declare that they have no competing interests.

**Correspondence and requests for materials** should be addressed to G.C.

## Notes

### Competing Interest Statement

The authors have declared no competing interest.

