## Supplementary information for "Spatial pattern and differential expression analysis with spatial transcriptomic data"

### Supplementary Methods

#### Estimation of the optimal hyperparameter $\theta_i$

In our main content, to better understand the construction of variance for the expression data $\boldsymbol{y}_i$, we express the total covariance as $\boldsymbol{\Sigma}_i = \gamma_{i1}\boldsymbol{K}_i + \gamma_{i2}\mathbf{I}$, which includes $\gamma_{i1}$ and $\gamma_{i2}$ to measure variance explained by spatial pattern and random noise, respectively. However, to simplify the derivation process for parameter estimation, we factor out a scaling variance factor $\gamma_i$ from $\boldsymbol{\Sigma}_i$, and redefine it as $\boldsymbol{\Sigma}_i = \gamma_i(\boldsymbol{K}_i + \varphi_i\mathbf{I})$, where $\varphi_i$ is a variance ratio factor with $\varphi_i = {}^{\gamma_{i2}}/_{\gamma_i}$. For simplicity of the presentation, we ignore the gene notation for all variables in the log likelihood in the following text (e.g., using $\boldsymbol{y}$ instead of $\boldsymbol{y}_i$).

$$\mathcal{L}(\boldsymbol{y}|\boldsymbol{X},\boldsymbol{\Theta}) = -\frac{N}{2}\log(2\cdot\pi) - \frac{1}{2}\log(|\boldsymbol{\Sigma}|) - \frac{1}{2}(\boldsymbol{y} - \boldsymbol{\mu}\cdot 1)^T(\boldsymbol{\Sigma})^{-1}(\boldsymbol{y} - \boldsymbol{\mu}\cdot 1),$$

where $\boldsymbol{\mu}$ is the mean gene expression level with $\boldsymbol{\mu} = \boldsymbol{X}^T\boldsymbol{\beta}$. Then we factor the kernel covariance matrix through spectral decomposition with $\boldsymbol{K} = \mathbf{VSV}^T$ to speed up the calculation of likelihood. Thus, the total variance can also be expressed as $\boldsymbol{\Sigma} = \gamma\cdot\mathbf{V}(\mathbf{S} + \varphi\cdot\mathbf{I})\mathbf{V}^T$, given that eigenvectors are orthogonal, $\mathbf{VV}^T = \mathbf{I}$. To estimate the mean expression level $\boldsymbol{\mu}$ and scaling variance $\gamma$, we take the first derivative of $\mathcal{L}(\boldsymbol{y}|\boldsymbol{X},\boldsymbol{\Theta})$ w.r.t. $\boldsymbol{\mu}$ and $\gamma$, respectively.

$$\frac{\partial\mathcal{L}(\boldsymbol{y}|\boldsymbol{X},\boldsymbol{\Theta})}{\partial\boldsymbol{\mu}} = 0 \rightarrow \widehat{\boldsymbol{\mu}} = \frac{(\mathbf{V}^T\mathbf{1})^T(\mathbf{S} + \varphi\cdot\mathbf{I})^{-1}(\mathbf{V}^T\boldsymbol{y})}{(\mathbf{V}^T\mathbf{1})^T(\mathbf{S} + \varphi\cdot\mathbf{I})^{-1}(\mathbf{V}^T\mathbf{1})}$$

$$\frac{\partial\mathcal{L}(\boldsymbol{y}|\boldsymbol{X},\boldsymbol{\Theta})}{\partial\gamma} = 0 \rightarrow \hat{\gamma} = \frac{1}{N}\cdot\frac{(\mathbf{V}^T\boldsymbol{y} - \mathbf{V}^T\mathbf{1}\cdot\widehat{\boldsymbol{\mu}})^T(\mathbf{V}^T\boldsymbol{y} - \mathbf{V}^T\mathbf{1}\cdot\widehat{\boldsymbol{\mu}})}{(\mathbf{S} + \varphi\cdot\mathbf{I})}$$

We found that the estimate of mean expression level $\widehat{\boldsymbol{\mu}}$ and scaling variance factor $\hat{\gamma}$ can both be expressed as a function depending only on $\varphi$, separately. Taking $\widehat{\boldsymbol{\mu}}$ and $\hat{\gamma}$ back to the log likelihood function $\mathcal{L}(\boldsymbol{y}|\boldsymbol{X}, \boldsymbol{\Theta})$, we can get an updated log likelihood function depending only on kernel hyperparameter $\theta$ and variance ratio factor $\varphi$. Thus, the optimal length-scale hyperparameter $\hat{\theta}$ and optimal $\hat{\varphi}$ can be simultaneously estimated by maximizing the log likelihood function $\mathcal{L}(\boldsymbol{y}|\boldsymbol{X}, \boldsymbol{\Theta})$. $\hat{\theta}$ will be used in the hypothesis test for identifying spatially variable genes.

### Supplementary Figures

**Figure S1: Distribution of estimated hyperparameter values in the simulation data with the MERFISH dataset.** Length-scale hyperparameter values for the pre-defined 50 marker genes were estimated using the SPADE method in the simulation data with the MERFISH data. Var: Variance.

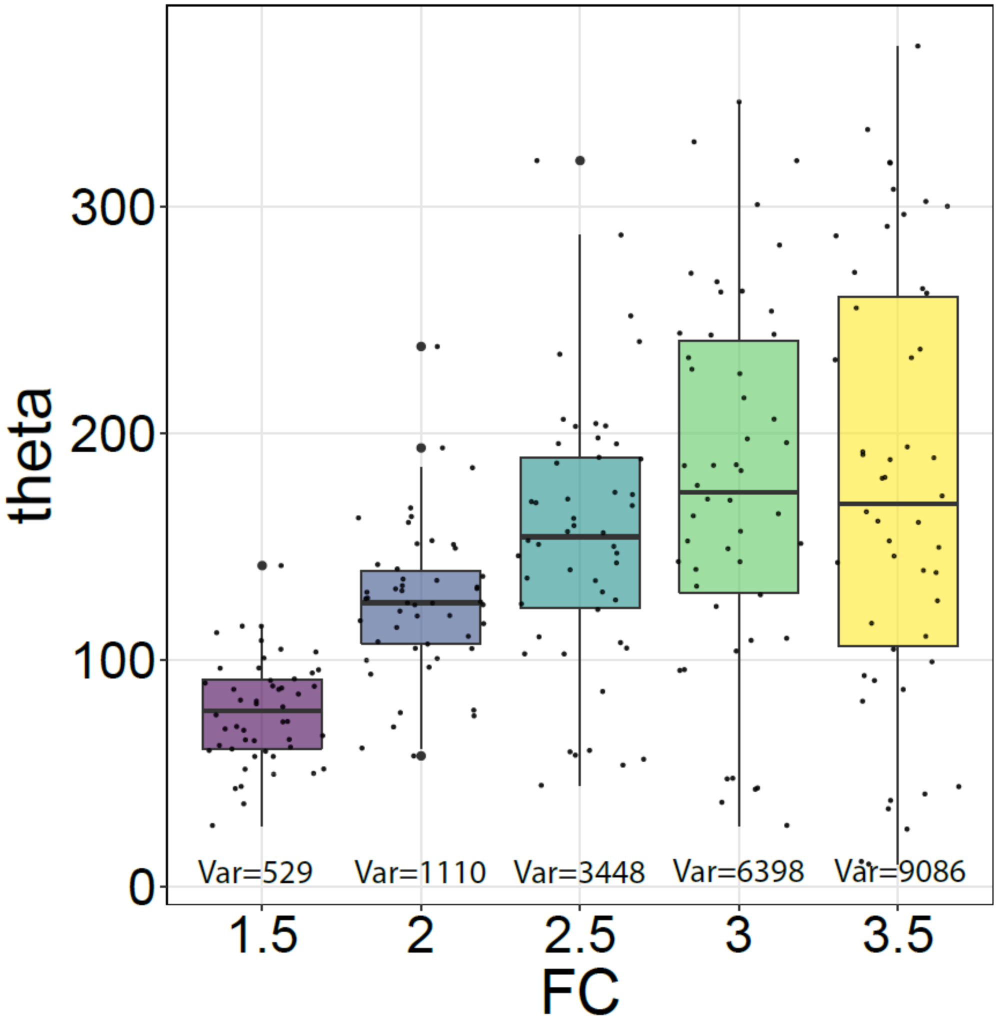

**Figure S2: Assessment of the robustness for SPADE to identify spatially variable genes between groups with marked spots proportion of 20% and hotspot patterns.** We evaluated the effects of spot coordinates and pattern directions to the performance of SPADE, separately. The areas under the receiver operating characteristic curve (AUC) were utilized to assess the power, and the false discovery rate (FDR) was used to assess the type I error of SPADE, respectively. Diff_coor: different coordinates; Same_coor: same coordinates; Diff_dir: different pattern directions.

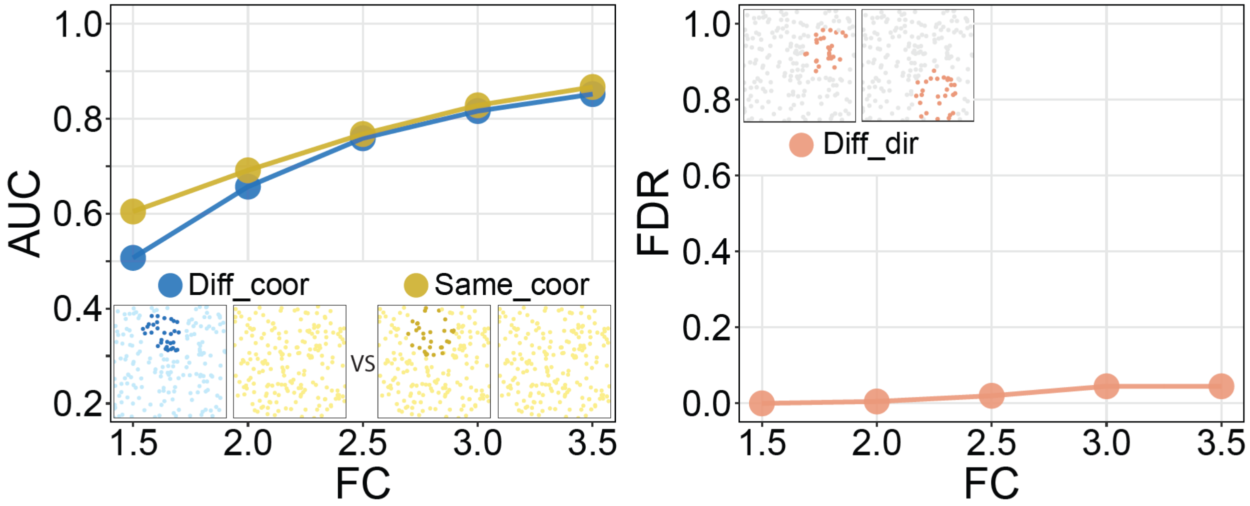

**Figure S3: Assessment of the robustness for SPADE to identify spatially variable genes between groups with marked spots proportion of 10%.** We evaluated the effects of spot coordinates and pattern directions to the performance of SPADE, separately. The areas under the receiver operating characteristic curve (AUC) were utilized to assess the power, and the false discovery rate (FDR) was used to assess the type I error of SPADE, respectively. Both streak (A) and hotspots (B) patterns were considered in simulation studies. Diff_coor: different coordinates; Same_coor: same coordinates; Diff_dir: different pattern directions; Same_dir: same pattern directions.

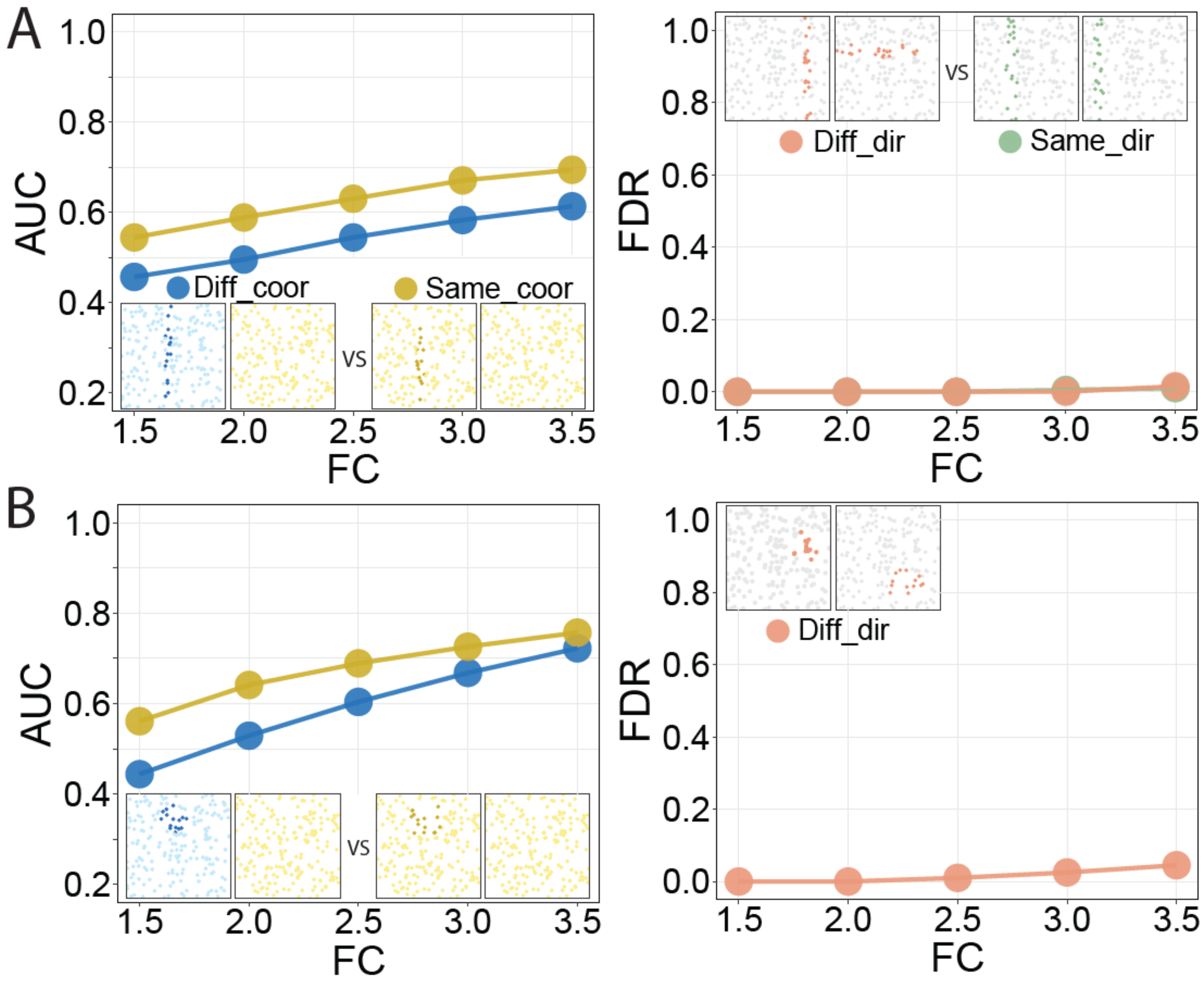

**Figure S4: Assessment of the robustness for SPADE to identify spatially variable genes between groups with marked spots proportion of 30%.** We evaluated the effects of spot coordinates and pattern directions to the performance of SPADE, separately. The areas under the receiver operating characteristic curve (AUC) were utilized to assess the power, and the false discovery rate (FDR) was used to assess the type I error of SPADE, respectively. Both streak (A) and hotspots (B) patterns were considered in simulation studies. Diff_coor: different coordinates; Same_coor: same coordinates; Diff_dir: different pattern directions; Same_dir: same pattern directions.

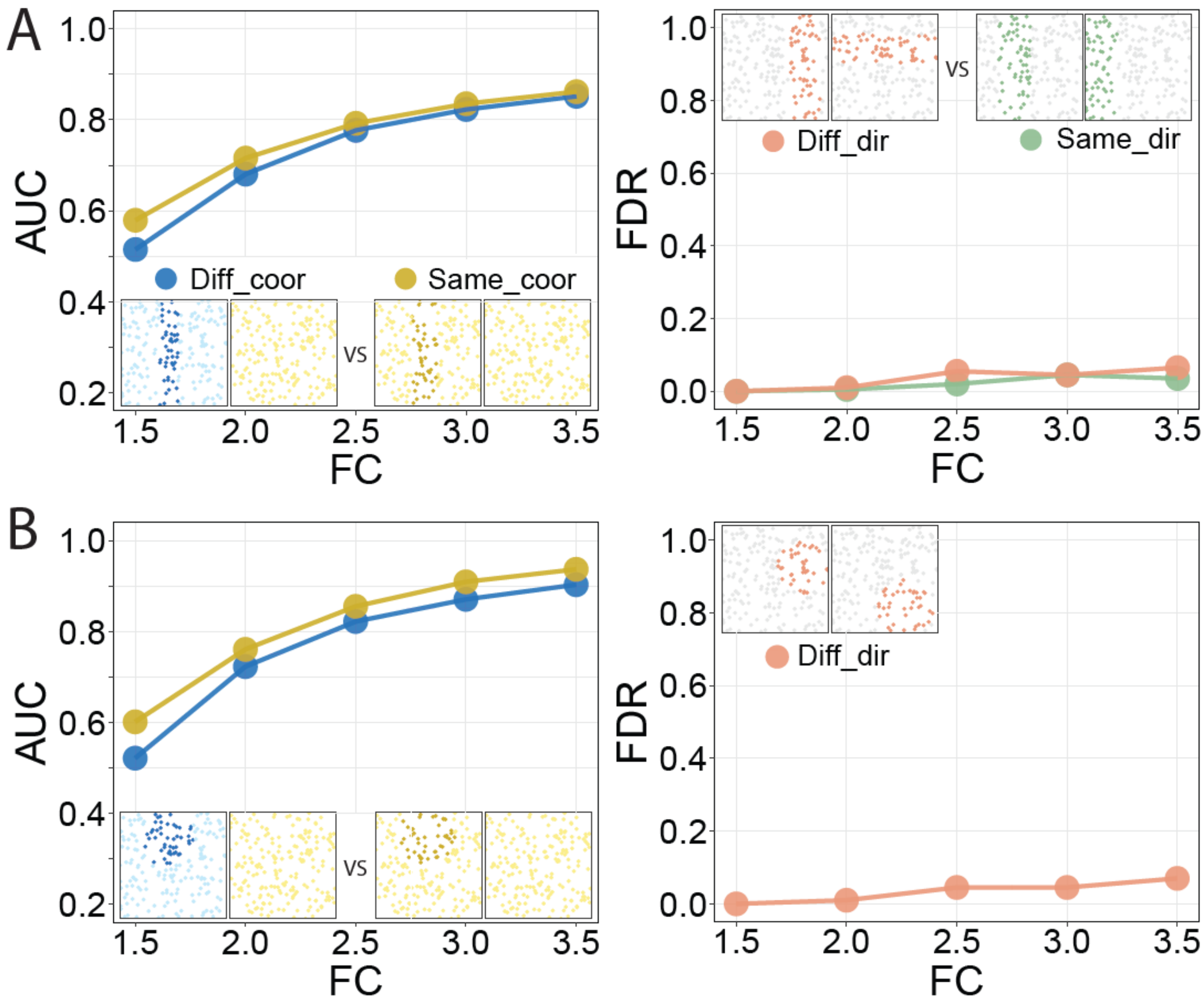

**Supplementary Table 1. Computational time of different methods with the SeqFISH dataset.** A desktop workstation with an Intel Core i5 CPU 2.6 GHz processor and 8.0 GB of RAM was used to identify spatially variable genes using SeqFISH dataset.

| Methods | Time (mins) |
|---|---|
| SPADE | 1.16 |
| SPARK | 1.08 |
| SpatialDE | 1.68 |
| MERINGUE | 0.01 |
